# Oxytocin Signaling Regulates the Homeostatic Response to Cold Stress in Poikilothermic Vertebrates

**DOI:** 10.1101/2021.12.20.472748

**Authors:** Adi Segev-Hadar, Shani Krispin, Anouk M. Olthof, Katery C. Hyatt, Liran Haller, Assaf Barki, Tali Nitzan, Gil Levkowitz, Rahul N. Kanadia, Avner Cnaani, Jakob Biran

**Author notes:** Correspondence should be addressed to J.B. These authors contributed equally to this work (Adi Segev-Hadar, Shani Krispin).

## Abstract

When exposed to low temperature, homeothermic vertebrates maintain internal body temperature by activating thermogenesis and by altered metabolism, synchronized by neuroendocrine responses. Although such physiological responses also occur in poikilothermic vertebrates, the prevailing notion is that their reactions are passive. Here, we explored molecular hypothalamic and physiological responses to cold stress in the tropical poikilotherm Nile tilapia (*Oreochromis niloticus*). We show that cold exposed tilapia exhibit complex homeostatic responses, including increased hypothalamic oxytocin, plasma glucose and cortisol concomitant with reduced plasma lactate and metabolic rate. Pharmacological or genetic blockage of oxytocin signaling further affected metabolic rate in two cold-exposed poikilothermic models. This indicates that oxytocin, a known thermoregulator in homeotherms, actively regulates temperature-related homeostasis in poikilotherms. Overall, our findings show that the brain of poikilotherms actively responds to cold temperature by regulating metabolic physiology. Moreover, we identify oxytocin signaling as an adaptive and evolutionarily conserved metabolic regulator of temperature-related homeostasis.

## Introduction

The notion of physiological homeostasis was conceived more than 140 years ago on the basis of equilibrium thermodynamics. The homeostatic view postulates that while biological systems are unstable by nature, they are regulated to maintain a dynamic equilibrium^1-3^. For example, temperature-related homeostasis is achieved in homeotherms through physiological and central mechanisms, such as metabolic heat generation. By contrast, in poikilotherms, commonly termed “cold-blooded” animals, it is achieved through behavioral modification, such as heat seeking behaviors in responses to cold stress^4^. Nevertheless, poikilotherms exposed to environmental extremes of cold temperatures experience a stressful metabolic challenge, which elicits physiological responses required to maintain cellular homeostasis. Moreover, heat seeking may not resolve the homeostatic needs of tropical poikilotherms under unpredictable extreme cold events, which occur frequently due to global climate change^5,6^.

Because poikilotherms cannot generate heat, it is expected that evolutionary pressure would lead their cells and tissues to develop responsive mechanisms to temperature alterations in order to maintain functionality and fitness under suboptimal conditions^7^. Indeed, previous studies have reported various responses to low temperature, including modified glucose or lipid metabolism^6,8-11^, altered gene expression and alternative splicing^10,12-14^, and endocrine and immune system activity^15,16^, which may vary according to the tissue and species. This suggests that the homeostatic response to cold stress in poikilotherms involves multiple physiological pathways and is more complex than currently perceived.

In vertebrates, the homeostatic activity of various organs and tissues is orchestrated by the brain hypothalamic region. The hypothalamus integrates sensory input from the internal and external environments and secretes regulatory neuropeptides and monoamines, which continually fine-tune physiological functions and maintain body homeostasis^17,18^. Several studies have demonstrated the thermoregulatory roles of hypothalamus in homeotherms^19,20^.

Bony fish (Class *Osteichthyes*), which comprise the largest group of vertebrates^21^, are poikilothermic^6,13^. As such, they have developed various strategies to survive in extreme cold waters, like hibernation and plasma antifreeze proteins. Yet, these strategies are utilized mainly by species inhabiting low-temperature environments, which encounter these conditions routinely throughout their life history^22^. In the last century, tropical fish species are increasingly exposed to extreme cold events due to several factors. First, climate change leads to increased weather events of extreme heat but also of extreme cold^5^. Second, the continuous expansion of the aquaculture industry leads to culture of tropical species in subtropical climates^23,24^. Third, aquaculture escapees, fish introductions for recreational angling and migration from saturated ecosystems result in species invasion into new ecosystems and ecological niches^23,25,26^. Exposure of a tropical fish to cold conditions would challenge its homeostasis and generate physiological stress. Yet, some species manage to survive exposure to cold temperatures and even thrive in climates that include cold seasons, which differ from their ecological life history. Understanding the central regulation of these homeostatic processes is important for ecological conservation and aquaculture, and may shed much needed light on the evolutionary origins of physiological thermoregulation in poikilothermic organisms.

Nile tilapia (*Oreochromis niloticus*) is one of the most important cultured fish worldwide. Originating from Africa, it is now cultured in more than 85 countries. With high reproductive rate, aggressive behavior, and a wide range of feeding sources, tilapia escapees have successfully invaded many ecosystems, including in sub-optimal climates^27,28^. Although some physiological and molecular homeostatic responses of tilapias to cold stress have been demonstrated^6,9,11,12,15^, the general metabolic relevance and hypothalamic regulation of these processes in poikilothermic fish remain mainly unexplored. In the present study, we examined the influence of cold exposure on Nile tilapia metabolism. As previously demonstrated in other piscine species^29^, we found a direct correlation between temperature and resting (standard) metabolic rate (RMR), as well as alterations in stress-related metabolic parameters upon exposure to extreme cold. In search of a central regulatory pathway that controls these physiological responses, we performed transcriptome analysis of Nile tilapia hypothalamus. Results showed that oxytocin (Oxt), a key regulator of core temperature in homeothermic mammals^30^, is markedly elevated upon exposure to extreme cold. Indeed, analysis of metabolic responses using pharmacological Oxt-receptor antagonist (ORA)^31,32^ in tilapia or genetic perturbation of Oxt signaling in zebrafish (*Danio rerio*) showed an Oxt-dependent decline in RMR during extreme cold exposure. These findings indicate that Oxt signaling is involved in the central thermoregulation of the physiological response to cold in poikilotherms, a function that was leveraged by homeotherms later in evolution. More broadly, our findings suggest that neuroendocrine pathways can modulate poikilothermic adaptiveness to climate change within physiological boundaries.

## Results

### Physiological response to cold stress

Survival temperatures of poikilothermic species are strongly correlated with their geographical distribution^33^, as reflected by the limited geographical expansion of Nile tilapia to tropical and subtropical areas^23,34^. Therefore, we utilized this fish as our model for studying regulation of homeostatic response to cold stress in poikilothermic vertebrates. To characterize the effects of cold temperature exposure on RMR and physiology of Nile tilapia, we analyzed the fish metabolic rate while reducing water temperature from 25°C to 14°C, at a rate of -1°C/h, followed by plasma analysis for major stress indicators and metabolic parameters (**Fig. 1**). This analysis showed direct correlation between temperature and RMR in Nile tilapia (**Fig. 1a-c**). In agreement with previous findings^11,15,35^, plasma cortisol and glucose levels significantly increased upon cold exposure, supporting a physiological stress response (**Fig. 1d-e**). Plasma lactate levels significantly decreased, which was also expected considering the reduction in RMR (**Fig. 1f**). These results are in line with cold-induced gluconeogenesis, which has previously been demonstrated by us and others^6,9^. No significant changes were found in plasma levels of total protein, triglycerides or growth hormone (**Fig. 1g-i**).

**Figure 1.**
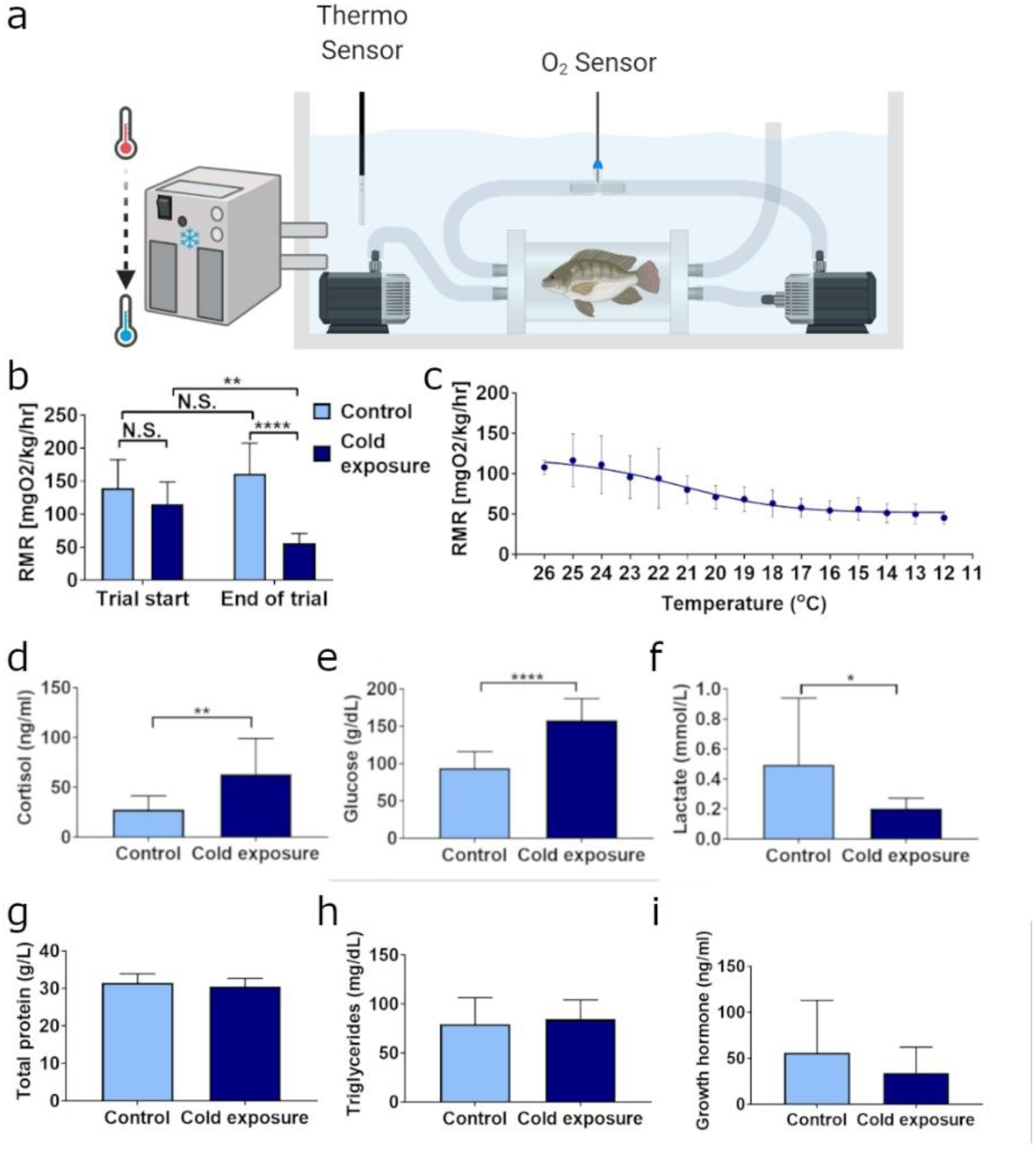
Cold exposure suppresses metabolic rate and elicits physiological stress response in Nile tilapia. (**a**) Schematic representation of the experiment. Fish were individually placed into 1L acrylic chambers connected to an oxygen measurement cell and submerged into temperature regulated water reservoir. (**b**) Analysis of the average metabolic rate at rest (RMR) of Nile tilapia prior to cold exposure (26°C and 25°C for control and cold-exposed fish, respectively) and following it (26°C and 14°C for control and cold-exposed fish, respectively) indicate a significant reduction in the RMR of exposed tilapia. (**c**) RMR analysis of cold-exposed fish in 1°C bins revealed a direct but nonlinear correlation of the fish RMR with the environmental temperature. (**d** and **e**) Plasma cortisol and blood glucose significantly increased following cold exposure; however, paradoxically, plasma lactate levels were decreased (**f**). (**g-i**) No significant changes were detected in plasma protein, triglyceride (TG) or growth hormone (GH) levels. The data are presented as mean ± SD.*p < 0.05; **p < 0.01; ****p < 0.0001. n=12 fish/treatment.

### Central pathways involved in the response to cold stress

In vertebrates, homeostatic functions are orchestrated by the hypothalamus, which serves as the central sensory and regulatory hub of peripheral body systems^17,36,37^. During an adaptive response, the hypothalamus generates a quiescent reaction to a strong metabolic challenge, while maintaining low physiological noise from other affected systems^3^. Hence, a reduction in the environmental temperature should induce a hypothalamic response that would change physiological and endocrine parameters, such as plasma levels of lactate, glucose and cortisol. To determine whether such homeostatic regulation occurs in fish hypothalamus, we dissected midbrains of Nile tilapia to include the diencephalon and optic tectum, which encompass all hypothalamic nuclei^38,39^ (**Fig. 2a**). Isolated midbrains from cold-exposed and normothermic fish were subsequently assessed for changes in gene expression levels by transcriptome analysis. First, the accuracy of midbrain dissections was confirmed by the expression of known hypothalamic neuroendocrine markers of the various hypothalamic regions, including *avp* (LOC100708704), *tac1, gal* (*galn*), *pomc* (*pomca*), *agrp* (LOC100691312), *trh, crh* (*crhb*) and others. Results showed that the expression of these genes was not significantly altered in the hypothalamus of cold-exposed Nile tilapia, compared to the controls (**Supp. Fig. 1**), confirming the integrity and uniformity of the hypothalamic dissections in both groups.

**Figure 2.**
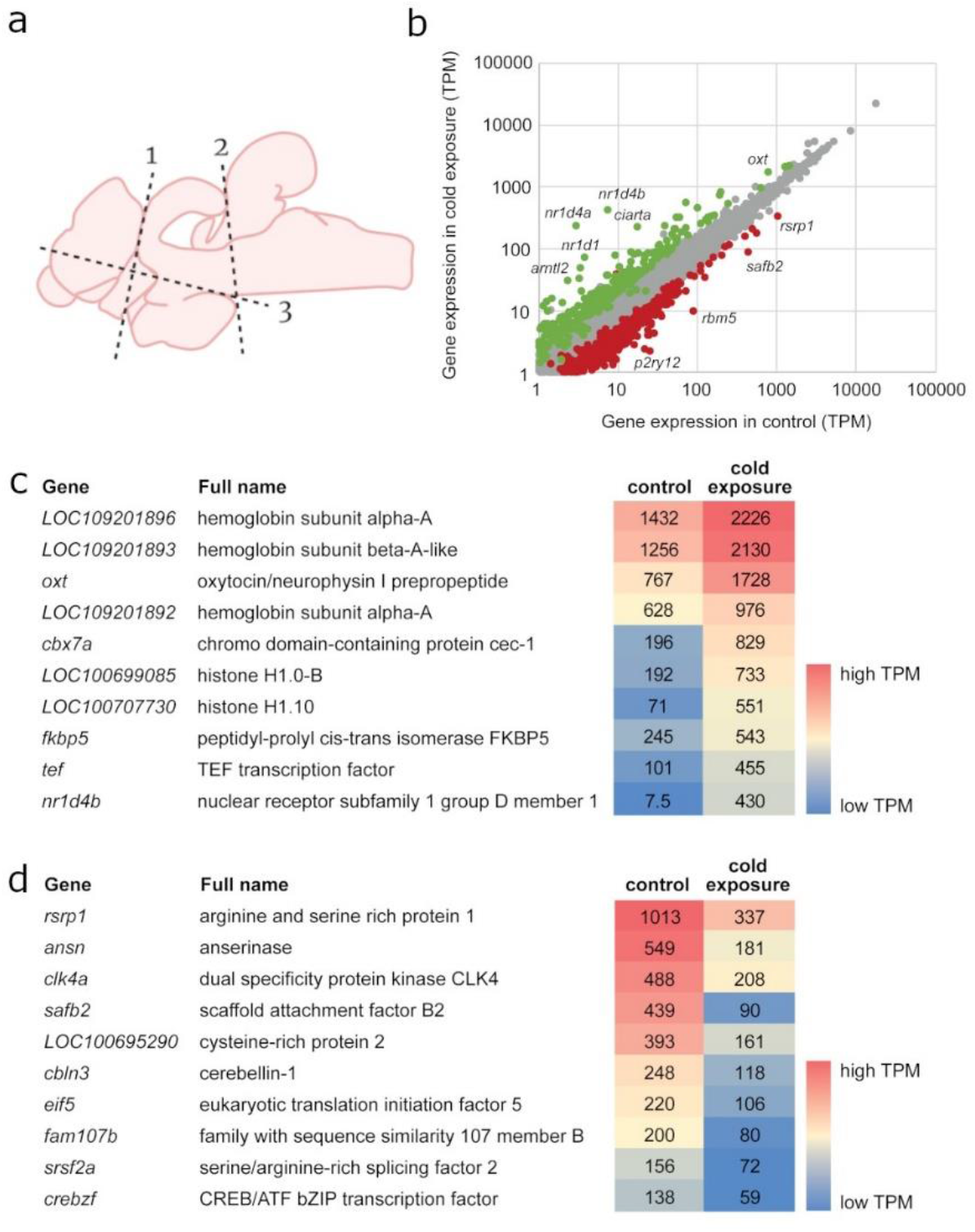
Cold exposure induces a transcriptional hypothalamic response of major neuroendocrine and metabolism related pathways. (**a**) Nile tilapia brains were micro-dissected to include all the major hypothalamic nuclei, including the preoptic area. (**b**) Transcriptome analysis of hypothalami revealed an increased expression of over 900 genes and suppressed expression of about 2,000 genes in response to cold exposure. (**c**) Analysis of the most highly expressed upregulated genes identified *oxt* as the most cold-responsive neuroendocrine factor, suggesting its involvement in the adaptive response of Nile tilapia to cold exposure. (**d**) Analysis of the most highly expressed downregulated genes yielded mainly genes involved in mRNA expression and processing. TPM, transcripts per million.

Next, we interrogated gene expression changes in the hypothalamus of cold-exposed Nile tilapia, using IsoDE2^40^. We found that 927 genes were significantly upregulated upon cold exposure, whereas 1971 genes were significantly downregulated (>2-fold change, P<0.01) (**Fig. 2b; Supp. Table 1**). To understand the biological processes that were affected by the differentially expressed genes, we next performed functional annotation analysis. Submission of the 911 downregulated genes that were expressed above 1 TPM (transcripts per million) in the control to g:Profiler^41^ yielded seven significant GO terms, which included broad functions such as “receptor signaling activity” and “peptide receptor activity” (**Table 1**). This suggested a generalized suppression of signaling pathways in Nile tilapia hypothalamus upon cold exposure. The 485 upregulated genes that were expressed above 1 TPM in the cold-exposed fish were significantly enriched for 21 GO terms (**Table 2**). These included the cellular component hemoglobin complex, apoptotic processes and cell death, as well as circadian rhythm pathways such as “circadian regulation of gene expression” and “rhythmic process”, suggesting that cold exposure affects the circadian rhythm of Nile tilapia (**Table 2**).

**Table 1.**
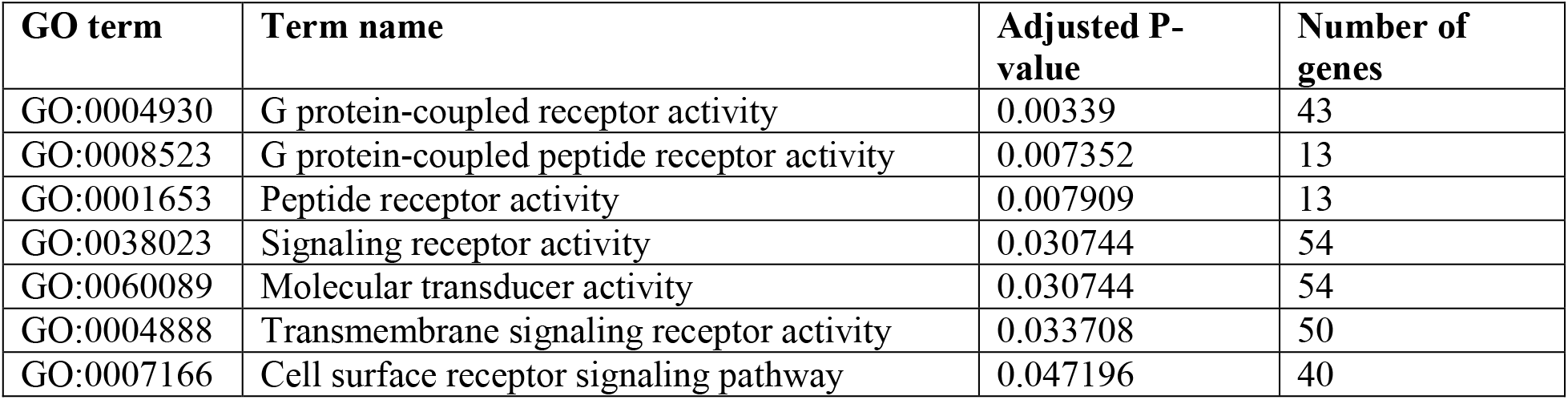
Functional annotation of genes downregulated upon cold exposure.

| **GO term** | **Term name** | **Adjusted P-value** | **Number of genes** |
| --- | --- | --- | --- |
| GO:0004930 | G protein-coupled receptor activity | 0.00339 | 43 |
| GO:0008523 | G protein-coupled peptide receptor activity | 0.007352 | 13 |
| GO:0001653 | Peptide receptor activity | 0.007909 | 13 |
| GO:0038023 | Signaling receptor activity | 0.030744 | 54 |
| GO:0060089 | Molecular transducer activity | 0.030744 | 54 |
| GO:0004888 | Transmembrane signaling receptor activity | 0.033708 | 50 |
| GO:0007166 | Cell surface receptor signaling pathway | 0.047196 | 40 |

**Table 2.** Functional annotation of genes upregulated upon cold exposure.

| GO term | Term name | Adjusted *P*-value | Number of genes |
| --- | --- | --- | --- |
| GO:0032922 | Circadian regulation of gene expression | 1.24E-6 | 6 |
| GO:0005634 | Nucleus | 3.50E-5 | 45 |
| GO:0003677 | DNA binding | 0.00024 | 37 |
| GO:0007623 | Circadian rhythm | 0.00031 | 7 |
| GO:0140110 | Transcription regulator activity | 0.00044 | 24 |
| GO:0048511 | Rhythmic process | 0.00046 | 7 |
| GO:0003700 | DNA-binding transcription factor activity | 0.00049 | 22 |
| GO:0046983 | Protein dimerization activity | 0.00493 | 16 |
| GO:0000981 | DNA-binding transcription factor activity, RNA polymerase II-specific | 0.00618 | 7 |
| GO:0006270 | DNA replication initiation | 0.01200 | 4 |
| GO:1901363 | Heterocyclic compound binding | 0.01401 | 90 |
| GO:0097159 | Organic cyclic compound binding | 0.01504 | 90 |
| GO:0097659 | Nucleic-acid templated transcription | 0.01522 | 36 |
| GO:0032774 | RNA biosynthetic process | 0.01596 | 36 |
| GO:0005833 | Hemoglobin complex | 0.01640 | 4 |
| GO:0098531 | Ligand-activated transcription factor activity | 0.01833 | 5 |
| GO:0004879 | Nuclear receptor activity | 0.01833 | 5 |
| GO:0043231 | Intracellular membrane-bounded organelle | 0.03371 | 49 |
| GO:0006915 | Apoptotic process | 0.03556 | 14 |
| GO:0012501 | Programmed cell death | 0.04078 | 14 |
| GO:0048523 | Negative regulation of cellular process | 0.04088 | 21 |
| GO:0003676 | Nucleic acid binding | 0.04799 | 58 |
| GO:0065007 | Biological replication | 0.04885 | 100 |

As expected by the altered metabolic rate, analysis of the most highly expressed upregulated genes revealed the presence of several genes encoding hemoglobin subunits, suggesting that the adaptive response of cold-exposed Nile tilapia may require increased brain oxygenation (**Fig. 2c**). While the general suppression of genes related to signaling pathways seems to support the concept of reduced metabolic responsiveness in cold-exposed poikilotherms^22^, we discovered that *oxt*, the gene encoding OXT neuropeptide, was significantly upregulated in the hypothalamus of Nile tilapia upon cold exposure (**Fig. 2c**). Similar analysis of the most highly expressed downregulated genes revealed suppression of several factors involved in the regulation of mRNA expression and processing (**Fig. 2d**). Our transcriptomic analysis was further validated by real-time PCR quantification of cold-induced (**Supp. Fig. 2**) and cold-suppressed (**Supp. Fig. 3**) mRNAs. These results support differential gene activation or suppression according to their involvement in specific cellular and physiological functions. Furthermore, the exceptional responsiveness of *oxt* expression to cold exposure suggests that OXT signaling is involved in the central regulation of the homeostatic response to cold stress in poikilotherms.

### OXT signaling reduces metabolism in poikilotherms under extreme cold conditions

The homeostatic response of homeothermic mammals to cold stress involves hypothalamic activation of OXT-neurons, which in turn elicit energy expenditure and thermogenesis to maintain core temperature^42,43^. However, in poikilothermic fishes low temperatures usually suppress feeding^44^ and, therefore, increased energetic expenditure may exhaust the energy storage of the animal and thereby risk its survival. Thus, we next aimed to determine whether the observed increase in *oxt* expression is related to poikilotherm thermoregulation through OXT signaling, or merely a result of globally altered regulation of gene expression. For this purpose, we used the OXT receptor-specific antagonist (ORA) L-368,899, which was shown to block OXT pathway from fish to mammals^31,32^. Nile tilapia were injected intraperitoneally with 1 mg/kg BW ORA and analyzed for their metabolic rate. Results showed that ORA significantly suppressed the cold-driven reduction in metabolic rate, an effect that was not detected in ORA-injected fish maintained in normothermy (**Fig. 3a-b**). Importantly, the effectiveness of ORA was seen only down to ∼19°C, suggesting that oxytocinergic regulation is only effective within the life history-shaped metabolic constrains of the species. This finding was accompanied by a significant reduction of plasma cortisol in ORA-treated fish exposed to cold stress (**Fig. 3c**). ORA did not affect plasma levels of glucose or lactate, supporting temperature-related metabolic rate regulation by OXT through modulation of physiological stress response.

**Figure 3.**
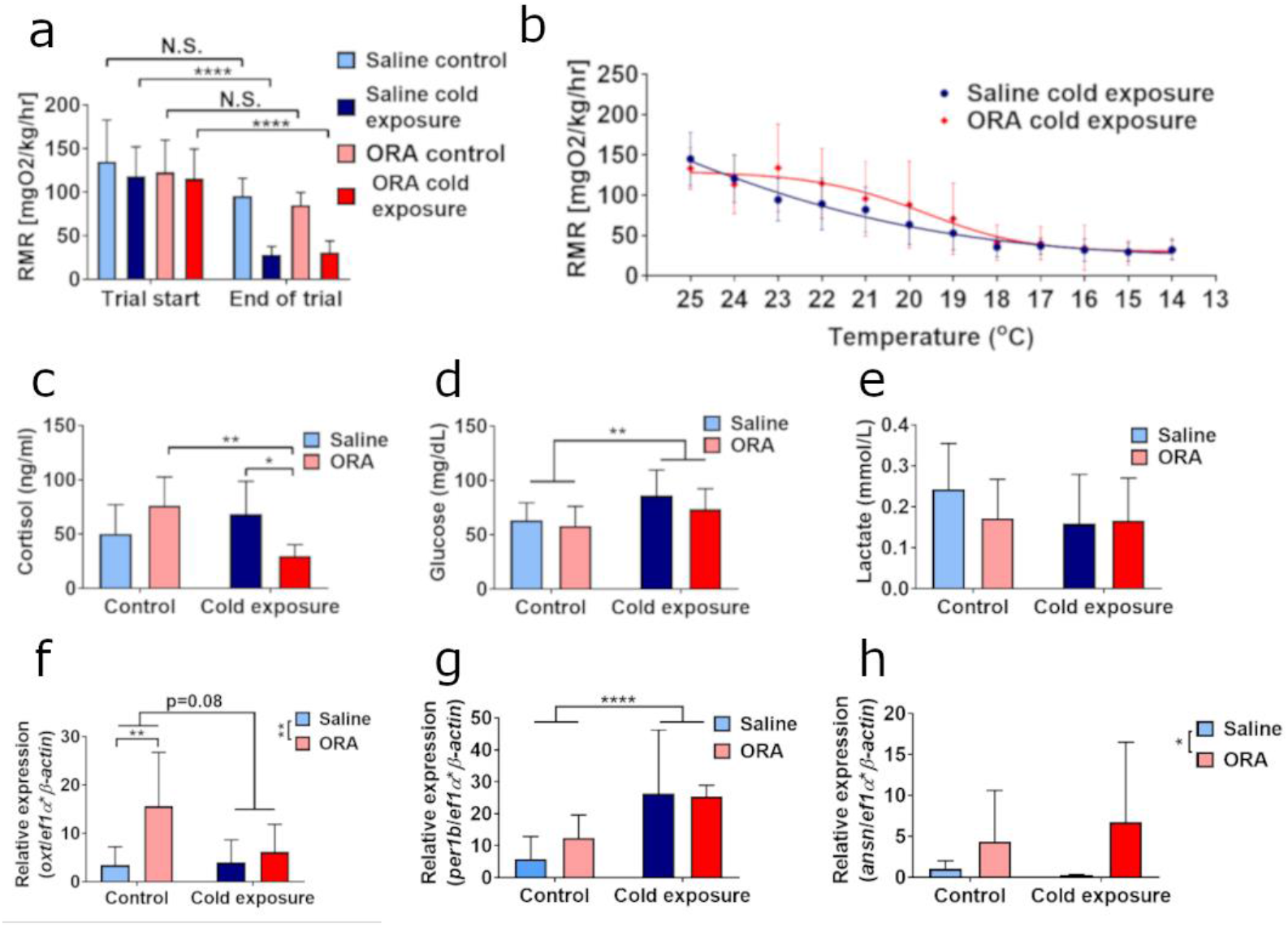
Oxt-receptor antagonist inhibits the cold-induced decline in metabolic rate and the physiological stress response in Nile tilapia. (**a**) Analysis of the average metabolic rate at rest (RMR) of Nile tilapia prior to cold exposure (24-24.5°C for all groups) and following it (24°C and 14°C for control and cold-exposed fish, respectively) indicate a significant reduction in the RMR of cold-exposed tilapia, regardless of the ORA treatment. (**b**) Nonetheless, regression analysis of normothermic and cold-exposed fish RMR in 1°C bins showed a significant perturbation of the fish RMR by ORA administration (p=0.0035). The cold-induced increase of plasma cortisol (**c**) but not of blood glucose (**d**) or lactate (**e**) was significantly blunted following ORA treatment. As expected by the blockage of OXT signaling, OXT expression was significantly increased by ORA (**f**). (**g**,**h**) While most analyzed genes were responsive to the cold exposure, the expression of the enzyme-coding *anserinase* responded significantly to ORA treatment. The data are presented as mean ± SD.*p < 0.05; **p < 0.01; ****p < 0.0001. n=12 fish/treatment.

The timeframe of the experiment was dictated by the previously reported pharmacokinetics of L-368,899^32^. Therefore, although ORA did not affect glucose and lactate levels (**Fig. 3d** and **Fig. 3e**, respectively), we could not exclude the involvement of OXT in gluconeogenesis and lactate metabolism, as these effects may require prolonged activation. ORA probably led to increased *oxt* expression due to activation of a feedback loop aimed to regain OXT receptor (OXTR) activity. Interestingly, this effect was more robust under normothermic conditions (**Fig. 3f**). Real-time PCR analysis of mRNAs which were also used for transcriptome validation showed that their expression was either unchanged or affected mainly by the change in temperature, rather than by ORA treatment (**Fig. 3g** and **Supp. Fig. 4**). The second most downregulated gene in response to cold, *anserinase* (*ansn*; **Fig. 2d**), is an orthologue of the homeothermic carnosinase enzyme unique to poikilothermic vertebrates^45,46^. Interestingly, ORA significantly induced *ansn* mRNA expression (**Fig. 3h**). These catabolic enzymes and their anserine/carnosine substrates have been associated with cognitive functioning, neurovascular activity and physiological homeostasis of histidine-containing dipeptides^45-47^. This further supports the specific activity of OXT in homeostatic regulation of tilapia’s metabolic response to cold stress.

### Evolutionary conservation of OXT signaling in temperature-related homeostasis

Our findings in Nile tilapia suggested that OXT signaling is a central regulatory pathway for temperature-related metabolic homeostasis in poikilotherms. To expand the evolutionary relevance of our findings, we used zebrafish (*Danio rerio*) as a complementary species. Although both species belong to the class of ray-finned fish (*Actinopterygii*), they are separated by over 300 million years of evolution^48^. Furthermore, while Nile tilapia originate from Africa, zebrafish originate from the Indian subcontinent and naturally experience a wider temperature range, making it more resilient to temperature extremes^9^. Thus, we used *oxt* and *oxtr* knockout (*oxt*^-/-^ and *oxtr*^-/-^, respectively) zebrafish germlines^49,50^ to analyze the involvement of OXT signaling in the central regulation of temperature-related metabolism in a distant poikilothermic species (**Fig. 4a)**. Our analysis demonstrated that under normothermic but not under extreme cold conditions, *oxtr*^-/-^ and, to a lesser extent, *oxt*^-/-^ mutant zebrafish display significantly reduced RMR (**Fig. 4b**). This finding suggests that OXT signaling is involved in maintaining basal metabolic functions in zebrafish. In view of the suppressed baseline RMR in *oxt*^-/-^ and *oxtr*^-/-^, the possible link between OXT signaling and zebrafish RMR during cold exposure was analyzed by subtracting the baseline RMR from the average RMR in each temperature (**Fig. 4c**). A direct correlation was found between reduced water temperature and zebrafish RMR; yet, this RMR suppression was faster in wild-type (WT) than in *oxt*^-/-^ and *oxtr*^-/-^ mutants (**Fig. 4c**). These findings support OXT signaling as a key pathway in the regulation of zebrafish baseline metabolic maintenance and cold-induced homeostatic adaptation.

**Figure 4.**
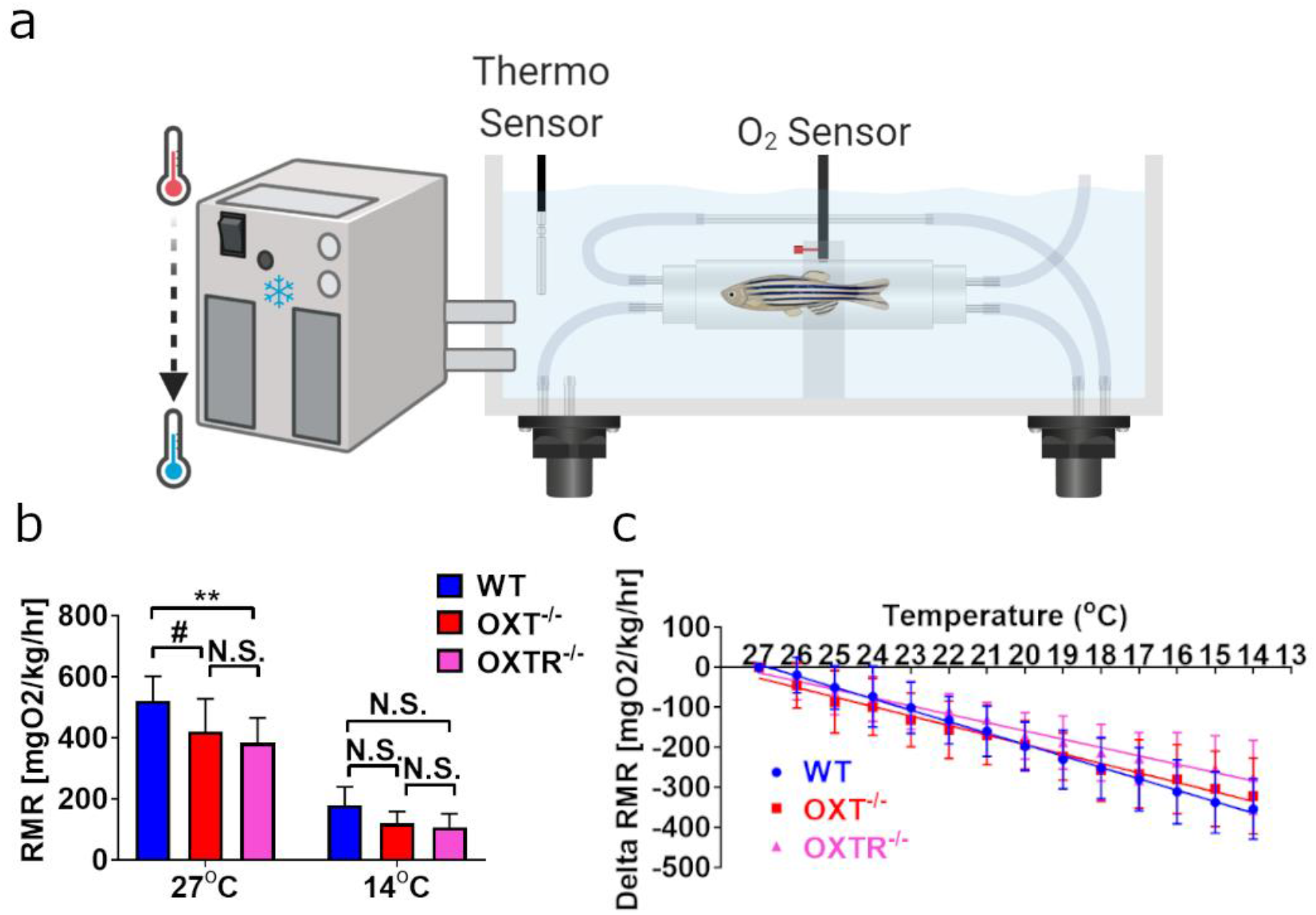
Genetic perturbation of OXT signaling alters basal and cold-adaptive metabolic rate in zebrafish. (**a**) Fish were individually placed into a glass chamber connected to an optical fiber for oxygen measurement, positioned in a temperature-regulated water reservoir. (**b**) As compared to wild-type (WT) controls, *oxt*^*-/-*^ and *oxtr*^*-/-*^ zebrafish displayed significant reduction in RMR under normothermic temperature (27°C), but not during cold exposure (14°C). (**c**) Therefore, delta RMR (RMR_temperature_-RMR_27oC_) was used to identify the adaptive cold-related effects of OXT perturbation, within the genetic background of each line. Similarly to the oxytocinergic effects detected in Nile tilapia, regression analysis of RMR data demonstrated that genetic ablation of *oxtr* or *oxt* significantly delayed RMR suppression caused by reduced water temperature (p=0.0013). The data are presented as mean ± SD. **p < 0.01; #p=0.0792; N.S., not significant. n=7-9 fish/treatment.

## Discussion

Homeostasis is a fundamental dogma in physiology. It states that environmental perturbations elicit physiological responses in the organism, which strive to regain stability and maintain fitness^1,3^. While the central and physiological responses of homeotherms to cold stress have been extensively studied, research of low temperature-related homeostasis in poikilothermic vertebrates has been narrowed to heat seeking behaviors, antifreeze protein production or hibernation^4,22^. Nevertheless, several studies demonstrated that active physiological and central modifications occur in poikilothermic fish exposed to cold stress^9,10,12,13^. Because the vertebrate hypothalamus serves as the homeostatic regulator of many physiological processes, its active response to extreme cold should support a thermoregulatory function. However, central pathways orchestrating such homeostatic responses have yet been identified^7^. Our current findings provide pioneering evidence for a central neuroendocrine regulation of metabolic rate in a poikilothermic vertebrate under cold stress conditions. We show that extreme cold exposure elicits a physiological stress response, which is accompanied by transcriptional upregulation of *oxt* in the midbrain-hypothalamic compartment. Next, we used an OXTR-specific antagonist and genetic KO models to demonstrate that OXT signaling regulates both metabolic rate and homeostatic physiology in cold-exposed poikilothermic vertebrates. Importantly, these data can also expand our understanding of the ecological impacts of globally increased incidences of extreme whether events and of invasive fish species, inadvertently introduced by the constantly expanding global aquaculture.

Acute cold exposure was shown to induce stress parameters including increased levels of plasma cortisol and catecholamines from fish to human^11,15,51^. However, while mammals exposed to extreme cold exhibit induction of energetic expenditure^4,51^, data by us and others^22^ show that rapid cold exposure in fish leads to reduced metabolic rate. As cold exposure suppresses feeding and activity in poikilothermic fish^44,52^, reduction in energetic expenditure is clearly beneficial for its fitness and survival under these conditions. Therefore, although these endocrine responses are widely conserved throughout evolution, their physiological manifestations should still differ according to the physiological constrains of the species and its life history.

The responses of poikilothermic fish to sharp declines in environmental temperatures were generalized to behavioral thermoregulation or cessation of physiological activity, whereas adaptive physiological mechanisms were considered only in polar or endothermic fish species^22^. Nevertheless, despite the expected RMR reduction in cold-exposed Nile tilapia, our data support an active homeostatic adaptation to the new conditions within the physiological boundaries of a tropical poikilotherm. This is well reflected by the increased levels of plasma cortisol, which is a known regulator of stress-related adaptation affecting multiple metabolic pathways^53^. Furthermore, cortisol was previously shown to significantly increase gluconeogenesis from lactate in the liver of several fish species^53^, which can explain why lactate levels remained low although glucose levels increased in cold-exposed fish. These data support a more active and complex homeostatic response to extreme cold than previously assumed.

Carbohydrate homeostasis is mainly performed in the liver^54^, corticosteroids are regulated by interrenal chromaffin cells of fish or adrenal cortex in mammals^17^, and catecholamine homeostasis involves synthesis in the autonomic nervous system and adrenal medulla and may also affect glucose homeostasis^55,56^. If these physiological changes are to increase the animal’s fitness, they must be synchronized with signals from the internal and external environments. While previous works have analyzed whole-brain transcriptomes^9,12,13^, our analysis focused in the vertebrates central homeostatic center, the hypothalamus^36^. Hence, in our search for a central regulator of the homeostatic response to cold stress in the poikilothermic Nile tilapia, we performed transcriptome analysis of the midbrain-hypothalamic compartment. The results suggested increased expression of genes involved in oxygen transport, cellular apoptosis and circadian rhythm pathways and suppressed expression of genes involved in pathways of peptide-related receptor signaling. This generally supports our initial hypothesis that poikilotherms exhibit active regulation of cold-driven homeostatic responses. Furthermore, the general suppression of peptide-related signaling is in line with our finding that cold exposure strongly affected midbrain *oxt* expression, suggesting that OXT is involved in the central regulation of the homeostatic response to cold stress. OXT is a known homeostasis-controlling neuropeptide involved in the regulation of metabolic physiology, behavioral and neuroendocrine stress responses and was recently suggested as a mediator of interactions between these homeostatic functions^30,57^. Additionally, OXT peptide sequence is evolutionarily conserved from worms to humans and so are some of its functions^58^. OXT and its cognate receptor are known regulators of mammalian core temperature by activation of physiological heat generation pathways^30,42^. OXT and OXTR mutant mice displayed impaired thermoregulation^42,59^ and their central recovery was sufficient to restore this function^43,60^, further supporting a direct involvement of OXT in temperature-related homeostasis. Nonetheless, to our knowledge, the role of OXT in temperature-related metabolic homeostasis of poikilothermic vertebrates has not yet been elucidated.

Strikingly, administration of ORA suppressed the temperature-dependent reduction in standard metabolic rate of cold-exposed tilapia. In addition, ORA affected plasma cortisol and central expression of specific mRNAs, supporting OXT signaling as an active and specific modulatory pathway of cold-related metabolic rate and physiology in a poikilothermic vertebrate model. These findings suggest that extreme cold exposure induce oxytocinergic signaling in the hypothalamus in order to suppress general energetic expenditure. Aiming to gain an evolutionary perspective of our findings, we analyzed our recently generated *oxt*^-/-49^ and *oxtr*^-/-50^ zebrafish lines under similar temperature challenge. Both mutant lines exhibited lower RMR under normothermy, suggesting that OXT signaling has an important role in maintaining general metabolism in the animal. This is in agreement with previous findings that mouse *Oxt*^-/-^ and *Oxtr*^-/-^ models demonstrate imbalanced energetic consumption versus expenditure^42^. Nonetheless, as seen in tilapia, mutant zebrafish exposed to extreme cold displayed lower RMR reduction rate compared to WT zebrafish.

In light of our current findings, we suggest that OXT signaling is a key regulator of low temperature-related metabolism in poikilothermic vertebrates. Moreover, we propose that instead of passive metabolism in poikilothermic vertebrates exposed to low temperatures^22^, there is an adaptive regulation of metabolic homeostasis, within physiological constrains. Therefore, while oxytocinergic signaling in homeotherms provoke internal heat production to maintain activity in cold environments, it actively suppresses energetic expenditure in cold exposed poikilotherms aiming to preserve energetic storage under low activity conditions. This notion should be incorporated into predictive modeling for aquaculture and invasive potential when considering introduction of non-native poikilotherms^23,25,28^, taking into account their homeostatic range in addition to their life history ecosystem. The relatively high amenability of aquaculture species to genome editing and rapid industry growth^61^ suggest that these considerations should also be applied to genome-edited lines intended for aquaculture in their native ecosystems. It was recently suggested that global temperature extremes may risk more than 60% of piscine species, mainly by affecting embryonal and reproductive life stages^62^. The high importance of OXT signaling to the embryonal development of hypothalamo-neurohypophyseal system^63^ and regulation of reproductive functions and behaviors were previously demonstrated from invertebrates to humans^58^. Thus, we suggest OXT as an evolutionarily conserved key neuroendocrine regulator of thermo-metabolic homeostasis in animals.

## Materials and Methods

### Animals

The experiments were approved by the Agricultural Research Organization Committee for Ethics in Using Experimental Animals (approval number: 806/18 IL). Nile tilapia males (body weight, 80±15 g) were raised in cylindrical 250-liter tanks (n = 8 fish/tank). Temperature was maintained at 24–26°C. Ammonia and nitrite levels were monitored. Fish were fed twice daily ad libitum with commercial tilapia feeds (Zemach Feed Mills™, Israel). Zebrafish (body weight, 0.49±0.12 g) were maintained under standard procedures as previously described^64^. Briefly, all genotypes were bred and reared at 28.5°C under 14 h/10 h light/dark cycle. Embryos were raised at 28.5°C in 30% Danieau’s medium supplemented with 0.01 mg/L methylene blue. Fish were deprived of food for 24 h prior to metabolic measurements to avoid digestion-related oxygen consumption.

### Metabolic rate analysis

The effect of gradual temperature decline on RMR was measured using an intermittent-flow respirometry system (Loligo Systems, Viborg, Denmark). Tilapia system included eight 1L acrylic cylindrical chambers that were equally allocated to control or cold exposure groups. Chambers were placed in a thermally regulated water tank with separated compartments of 77 L each. Temperature of control group was maintained at 26°C (±0.5°C) using a standard heating element, while temperature of treatment group was regulated using a refrigerated/heated bath circulator (Arctic Series A25, ThermoFisher Scientific, USA) linked to a submerged heat exchanger coil. Oxygen levels in tanks were maintained at 90%-100% O_2_ saturation using multiple air stones and water were recirculated through a UV lamp apparatus to avoid bacterial growth. Each respirometric chamber was connected to two separate water pumps (Eheim, Germany). One was used for flushing between subsequent measurements, whereas the other was used for recirculating water to allow for dissolved oxygen measurement via a flow-through oxygen cell and a mini spot sensor connected through an optical fiber to a Witrox oxygen meter. Temperature was continuously monitored using a software-integrated thermometer; system operation as well as data monitoring were performed using AutoResp software (ver. 2.2.2, Loligo Systems). The respirometric system was drained and cleaned between runs to prevent the development of biofilm which may cause background respiration.

The experiment included 24 fish (n=12 fish/treatment) and was divided into 3 subsequent and identical runs of 8 fish per run (n=4 fish/treatment) over the course of one week. In each run, fish were weighed and randomly placed in either treatment or control chambers for a 4-hour acclimation. Overnight measurements began at 5 pm and lasted for 15 h, during which temperature for treatment group was reduced from 26°C to 14°C (±0.5°C) at an average rate of 0.75°C/h. Mass-specific O_2_ consumption (ṀO_2_; mg O_2_/kg/h) was continuously measured in 8.5 minute cycles, each composed of a “flush” (3 min), “wait” (0.5 min) and “measure” (5 min) periods. Duration of measurement cycles was empirically tested to avoid reaching below 80% O_2_ saturation and minimize additional effect of respiratory stress. For each animal, RMR was analyzed in 1°C (±0.5) bins.

Zebrafish system included eight 11.5 mL glass cylindrical chambers. Chambers were placed in one of two thermally regulated 10 L water tanks connected through a 70 L reservoir. Temperature was maintained at 27°C (±0.5°C) using a standard heating element followed by gradual decrease averaged at ∼1°C/h using a refrigerated/heated bath circulator (Arctic Series A25, ThermoFisher Scientific, USA) linked to a heat exchanger coil submerged in the main reservoir. Oxygen levels in tanks were maintained at 90%-100% O_2_ saturation. Each respirometric chamber was connected to two separate miniature impeller pumps (PU10700, Loligo Systems). Flow scheme and regulation were as described for tilapia. ṀO_2_ was continuously measured using 300 seconds cycles, each composed of a “flush” (90 seconds), “wait” (30 seconds) and “measure” (180 seconds) periods. To avoid inter-measurements bias, every trial contained 1-2 fish from each genotype, which were randomly assigned to the respirometric chambers. For each animal, RMR was analyzed in 1°C (±0.5) bins.

### Pharmacological treatment

To assess the involvement of OXT signaling in the metabolic rate at rest of cold-exposed Nile tilapia, fish were intraperitoneally injected with L-368,899 (ChemCruz, Dallas, TX, USA), which is a known ORA^31,32^. Control fish were intraperitoneally injected with saline. Metabolic analysis of pharmacologically treated fish was performed as described above, with slight modifications. Analysis was performed in four replicates, each included 4 control (2 normothermy and 2 cold-exposed) and 4 ORA-treated (2 normothermy and 2 cold-exposed) fish. In view of the previously described pharmacokinetics of L-368,899^29,30^, temperature reduction in the cold-exposed group was modified to 1.25-1.5 °C/h and started immediately following ORA administration.

### Tissue collection and biochemical analysis

At the end of each run, blood was collected via the caudal vein using a 23-gauge hypodermic needle rinsed with heparin (200 IU/ml). Blood glucose levels were measured using a FreeStyle Optimum glucometer (Abbott Diabetes Care, Witney, UK). Plasmas were separated from blood cells and platelets by centrifugation at 4°C/3.2 g for 20 min, transferred to 1.5 mL tubes and stored at -80°C until further analysis. Following blood collection, weight and total length were measured for each fish and the diencephalon, including the preoptic area, were micro-dissected and snap-frozen in liquid nitrogen. Subsequently, plasma lactate, triglycerides and total protein content, as well as cortisol and growth hormone (GH) levels, were measured. Lactate, triglycerides and total protein were quantified using the Cobas c111 analyzer (Roche diagnostics GmbH, Mannheim Germany) as previously described by Segev-Hadar et al.^38^. Cortisol and GH were quantified according to previously published protocols by Yeh et al.^65^ and Mizrahi et al.^66^.

### RNA extraction and transcriptome sequencing

Total RNA was extracted using TRIzol® reagent (Life Technologies Corporation, Carlsbad, USA) according to manufacturer protocol and purified to remove remaining DNA contamination using the TURBO DNA-free™ kit (Invitrogen, USA). RNA concentration and purity were determined using an Epoch Microplate Spectrophotometer (BioTek, USA). RNA samples from treatment and control groups (n=4/treatment) were sequenced at the Technion Genome Center (Technion Institute of Technology, Haifa, Israel). RNA integrity was tested using an Agilent 2200 TapeStation (Agilent Technologies, USA). Subsequently, poly (A) mRNA was isolated from the total RNA with poly (dT) oligo-attached magnetic beads, and cDNA libraries were prepared using the TruSeq RNA Sample Preparation Kit (Illumina, USA) following the manufacturer protocol. Eight cDNA libraries were sequenced on a single lane by the HiSeq2500 sequencing platform (Illumina, USA) at 2 × 100 bp paired-end (PE) reads. The data have been deposited in the GEO database (accession number GSE159019).

### Gene expression analysis

The *O. niloticus* genome was downloaded from NCBI on April 2019 (v.1.0). Reads were aligned to the genome using Hisat2^67^, and gene expression was determined using IsoEM2. Differential gene expression was determined using IsoDE2^40^, which reports a confident fold-change (*P*<0.01). Genes with a confident fold-change >2 were considered differentially expressed. Upregulated and downregulated genes expressed above 1 TPM in at least one condition were submitted to g:Profiler^41^ for functional annotation analysis using *O. niloticus* as background. Real-Time PCR validation of selected genes was performed as previously described^38^. Briefly, possible genomic DNA contamination was eliminated by treatment with Invitrogen TURBO DNA-freeTM kit (Thermo Fisher Scientific, Vilnius, Lithuania) according to the manufacturer’s protocol. DNase-free total RNA (0.5 μg) was reverse-transcribed using Verso cDNA kit (Thermo Fisher Scientific; naïve fish) or High Capacity cDNA Reverse Transcription kit (Thermo Fisher Scientific; ORA experiment) according to the manufacturer’s protocol. cDNA was stored at −20°C until quantitation by real-time PCR. Hypothalamic gene expression levels were analyzed by quantitative PCR using a StepOnePlus Real-Time PCR System (Applied Biosystems, Inc. Foster City, CA, USA). *elongation factor 1 alpha* (*ef1α*) and *β-actin* served as reference genes (**Supp. Table 2**). Each reaction consisted of 5 μL SYBRR green dye (Thermo Fisher Scientific, Vilnius, Lithuania), 0.5 μL of 2 μM forward and reverse primers, 2.5 μL of ultra-pure water (UPW) and 1.5 μL of cDNA template (diluted 1:16 in UPW). Analysis was performed in duplicates. Controls without the cDNA were used to test for non-specific amplification. Specificity of the primers was validated by Sanger sequencing and melt curve analysis was used to confirm amplification of a single product. Amplification was performed under the following conditions; 95.0°C for 20 sec, 40 cycles at 95.0°C for 3 sec, and 60.0°C for 30 sec, followed by one cycle at 95.0°C for 15 sec and 60.0°C for 1 min, 95.0°C for 15 sec for the generation of the melting curve. Fluorescence signals of the target and reference genes in the control and treatment groups were analyzed using StepOne software Version 2.3. Relative quantification of within-tissue expression was determined using the 2^-ΔΔ^CT method^68^.

### Statistical analysis

Values are presented as mean ± standard deviation (SD). The level of significance was set to p < 0.05 in all performed analyses. A two-way ANOVA (Tukey’s multiple comparisons) was used to test for RMR differences between normothermic and cold-exposed fish. Unpaired Student’s t-test test was used for analyzing the physiological parameters measured and for analyzing Real-Time PCR data of transcriptome validation. Analysis of RMR differences of the pharmacological analysis demonstrated that the data significantly diverged from linearity. Therefore, the data were analyzed using non-linear regression followed by an extra sum-of-squares F test which demonstrated that the data sets could not be represented by a single curve (p=0.0035). Analysis of physiological parameters measured and Real-Time PCR data were performed using a two-way ANOVA (Tukey’s multiple comparisons). A two-way ANOVA (Tukey’s multiple comparisons) was used to test for RMR differences between different zebrafish genotypes at specific temperatures. Due to baseline RMR differences of the zebrafish mutants, data were plotted as delta RMR (RMR[27°C]-RMR[X°C]). Data sets were further analyzed by linear regression which demonstrated that slopes are significantly different (p=0.0013). Statistical analyses performed using GraphPad Prism 7.03.

## Acknowledgements

We thank Tatiana Slosman (Agricultural Research Organization) and Roy Hofi (Weizmann Institute of Science) for animal care and Nitzan Konstantin for English editing. This research was supported by grants 20-04-0055 (to J.B.) and 20-11-0026 (to A.C.) from the Chief Scientist of the Ministry of Agriculture and Rural Development. We thank Jannik Herskin and Andreas Mørck (Loligo Systems) for their technical assistance and graphical contribution of metabolic systems to Figures 1a and 4a. Other components in Figures 1a and 4a were created with BioRender.com.

## Author contributions

J.B., A.C. and R.N.K. conceived and designed the project. A.S.H., S.K., L.H., A.B. and T.N. performed *in vivo* metabolic analyses, physiological and molecular analyses. A.M.O, K.C.H and R.N.K performed the bioinformatics analysis. G.L. designed the metabolic analysis of zebrafish mutants and contributed *oxt*^-/-^ and *oxtr*^-/-^ germlines. J.B. prepared the figures. J.B., A.C., R.N.K. and A.M.O. wrote the manuscript. All authors reviewed the manuscript.

## Supplemental Information

**Supplemental Figure 1.**
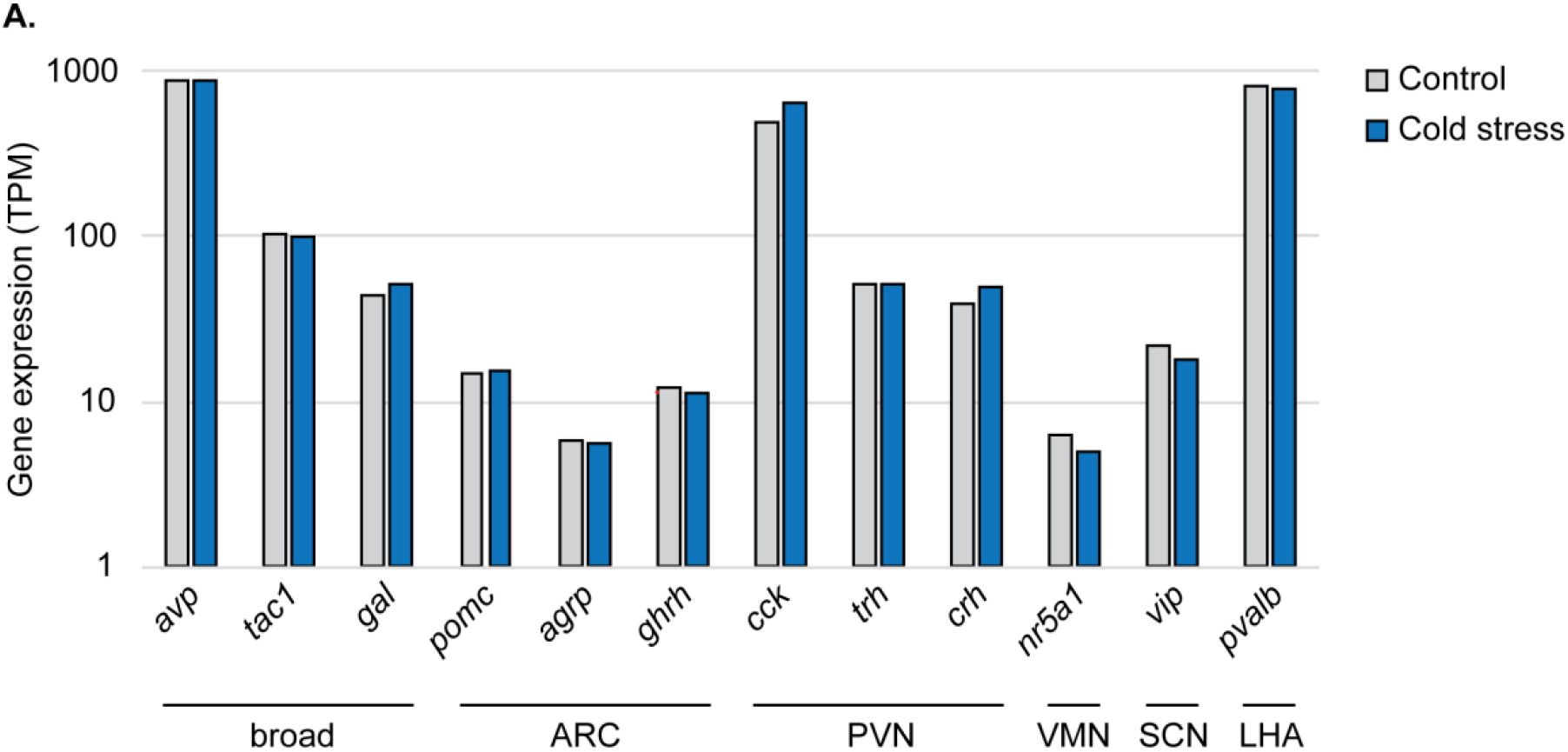
Validation of hypothalamic microdissection by RNAseq analysis for expression of hypothalamic markers. No significant differences were observed for *LOC10070874* (*avp*), *tac1, gal, pomc* (*pomca*), *LOC100691312* (*agrp*), *ghrh, cck, trh, crh* (*crhb*), *nr5a1, LOC100705021* (*vip*) or *LOV100710987* (*pvalb*). Genes are grouped by the hypothalamic nucleus they mark in the mammalian brain. ARC, arcuate nucleus; PVN, paraventricular nucleus; VMN, ventromedial nucleus; SCN, suprachiasmatic nucleus; LHA, lateral hypothalamic area.

**Supplemental Figure 2.**
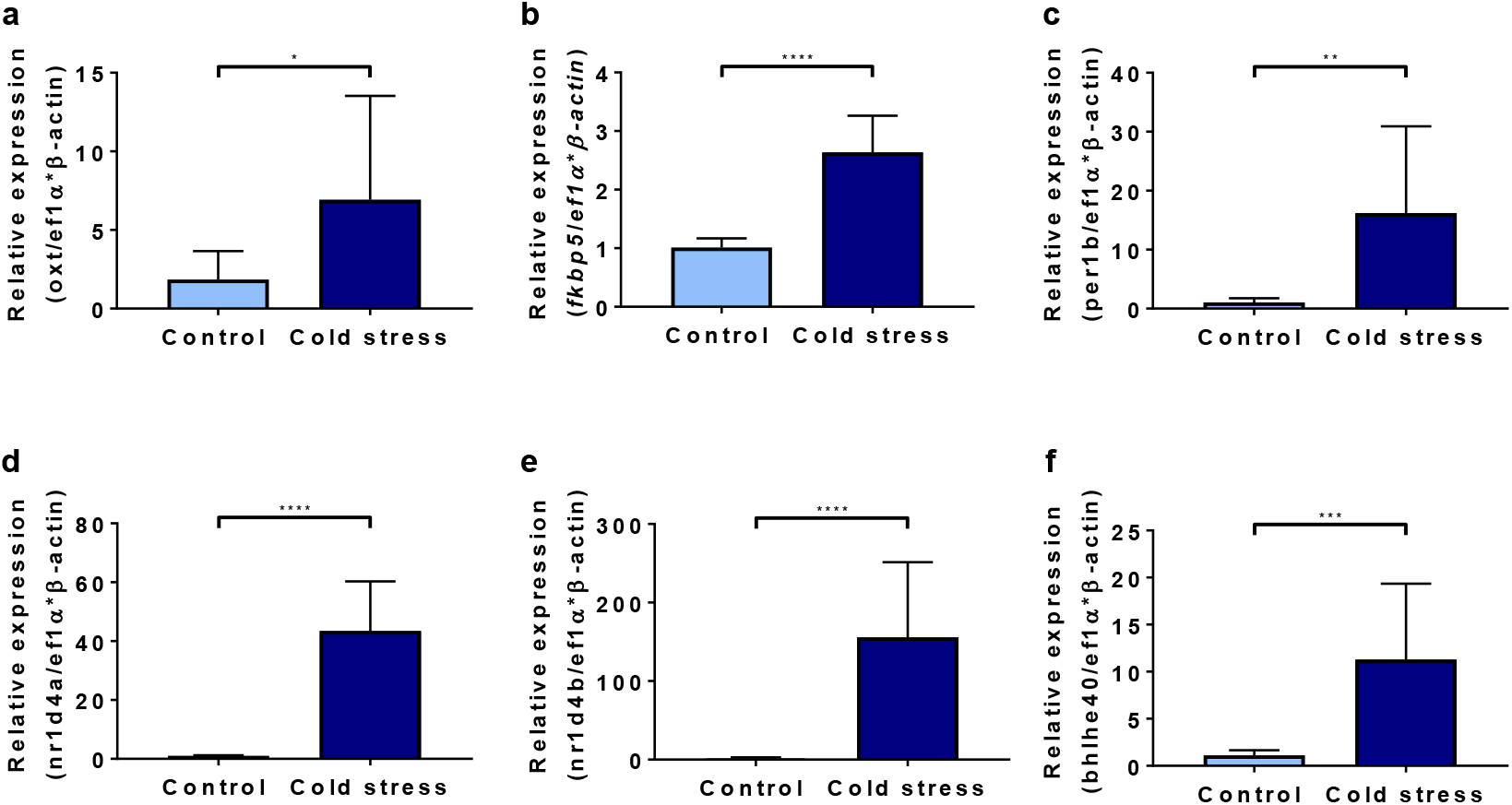
Validation of transcriptomic analysis results for identified cold-induced genes. Total RNA was extracted from midbrains of control (n=11) and cold-exposed (n=9) tilapia and was analyzed by real-time PCR for cold-induced expression of various genes. Similar to the transcriptome data, *oxytocin* (**a**), *fkbp5*(**b**), *per1b* (**c**), *nr1d4a* (**d**), *nr1d4b* (**e**) and *bhlhe40* (**f**) displayed significantly increased expression in response to cold exposure. The data are presented as mean ± SD.*p < 0.05; **p < 0.01; ***p < 0.001; ****p < 0.0001.

**Supplemental Figure 3.**
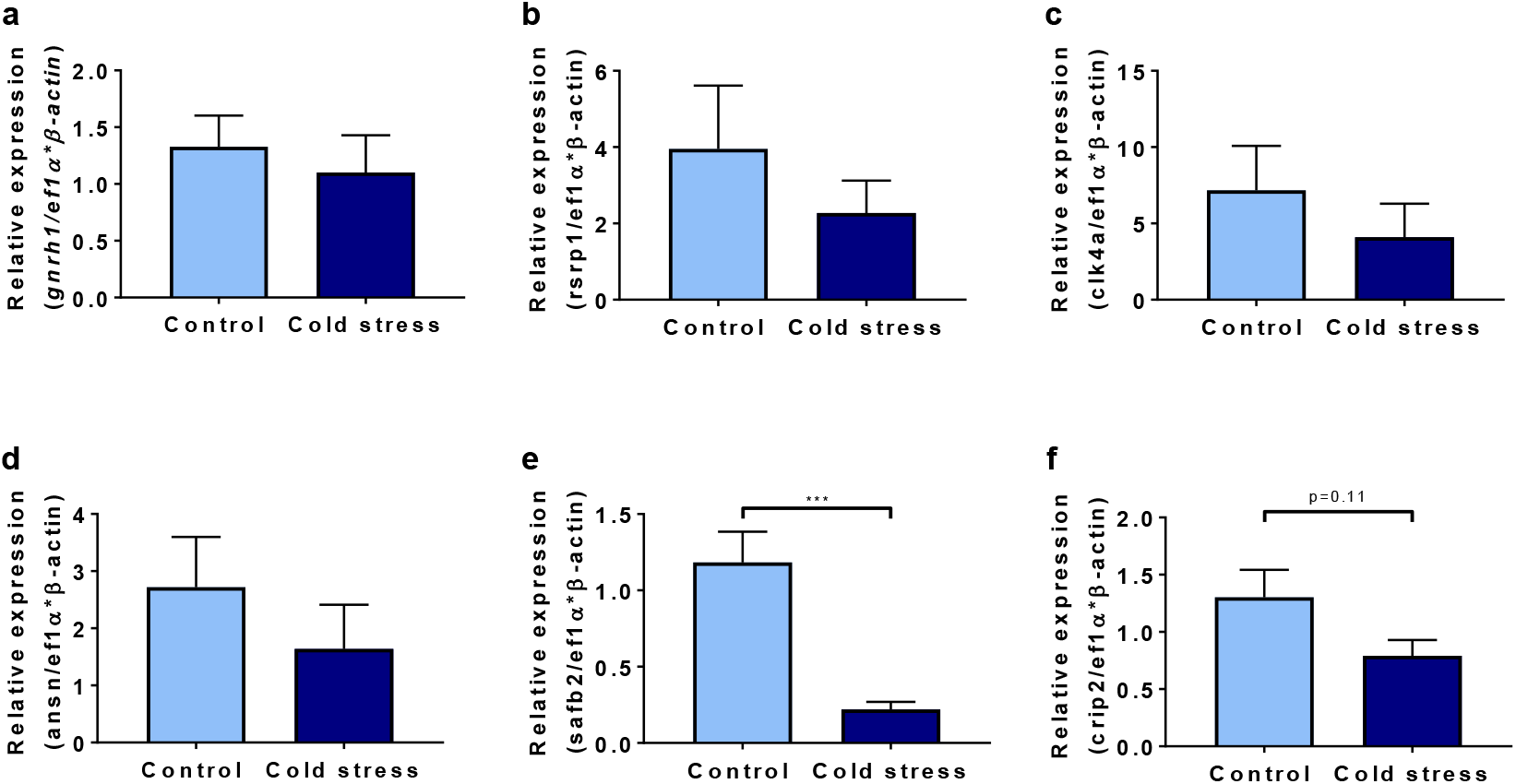
Validation of transcriptomic analysis results for identified cold-suppressed genes. Total RNA was extracted from midbrains of control (n=11) and cold-exposed (n=9) tilapia and was analyzed by real-time PCR for cold-suppressed expression of various genes. Similar to the transcriptome data, *gnrh1* (**a**), *rsrp1* (**b**), *clk4a* (**c**), *ansn* (**d**), *safb2* (**e**) and *crip2* (**f**) displayed clear trends for decreased expression in response to cold exposure. The data are presented as mean ± SD. ***p < 0.001.

**Supplemental Figure 4.**
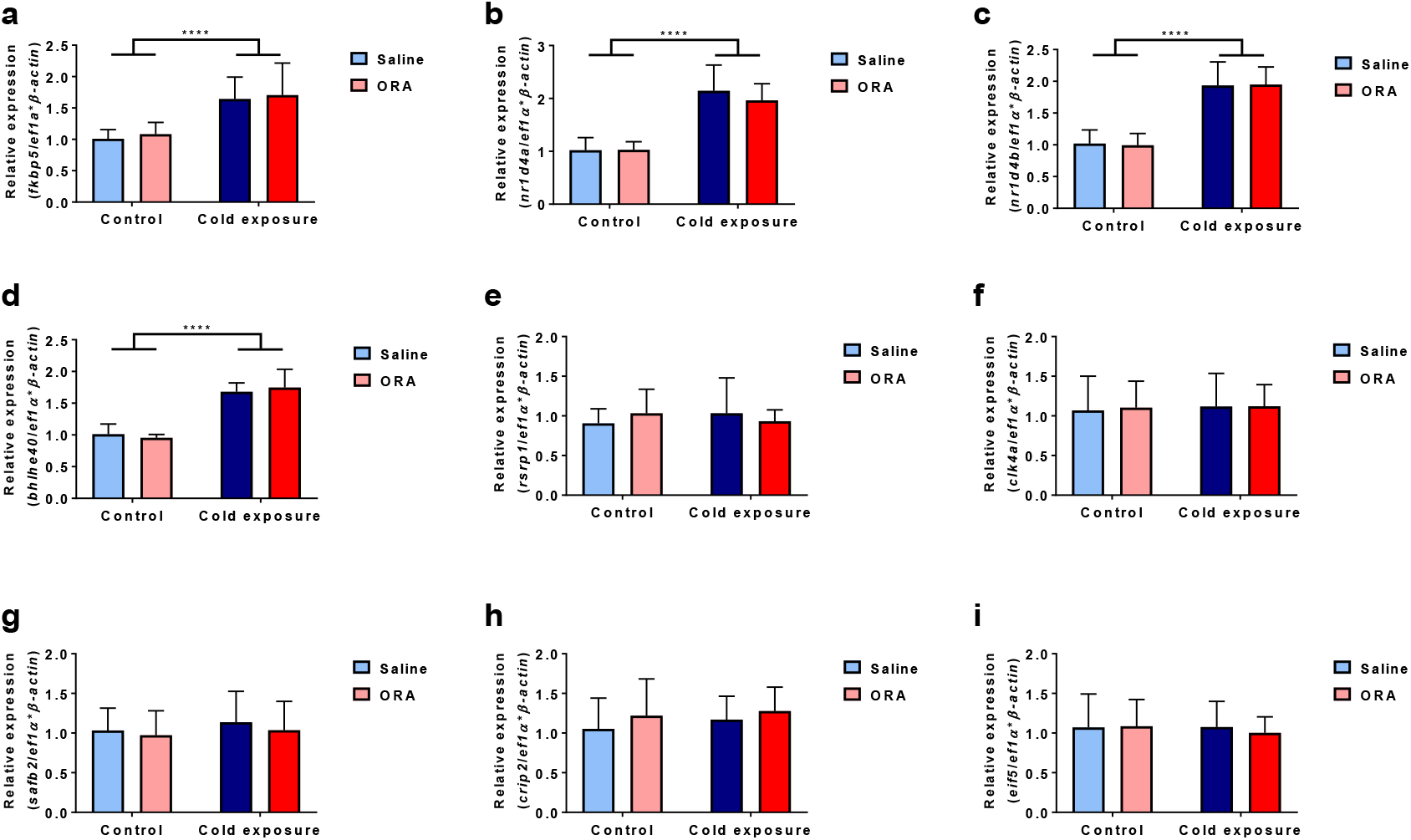
Expression analysis of cold-responsive genes upon administration of Oxt-receptor antagonist. Total RNA was extracted from midbrains of control and cold-exposed tilapia that were intraperitoneally injected with either saline or 1 mg/kg body weight ORA and was analyzed using real-time PCR. Most genes displayed similar trends of responsiveness as we identified in non-injected tilapia. Analyzed genes included *fkbp5* (**a**), *nr1d4a* (**b**), *nr1d4b* (**c**), *bhlhe40* (**d**), *rsrp1* (**e**), *clk4a* (**f**), *safb2* (**g**), *crip2* (**h**) and *eif5* (**i**). The data are presented as mean ± SD. ****p < 0.0001. n=8 fish/treatment.

**Supplemental Table 1. Gene expression of cold-exposed Nile tilapia**

See attached Excel file

**Supplemental Table 2.**
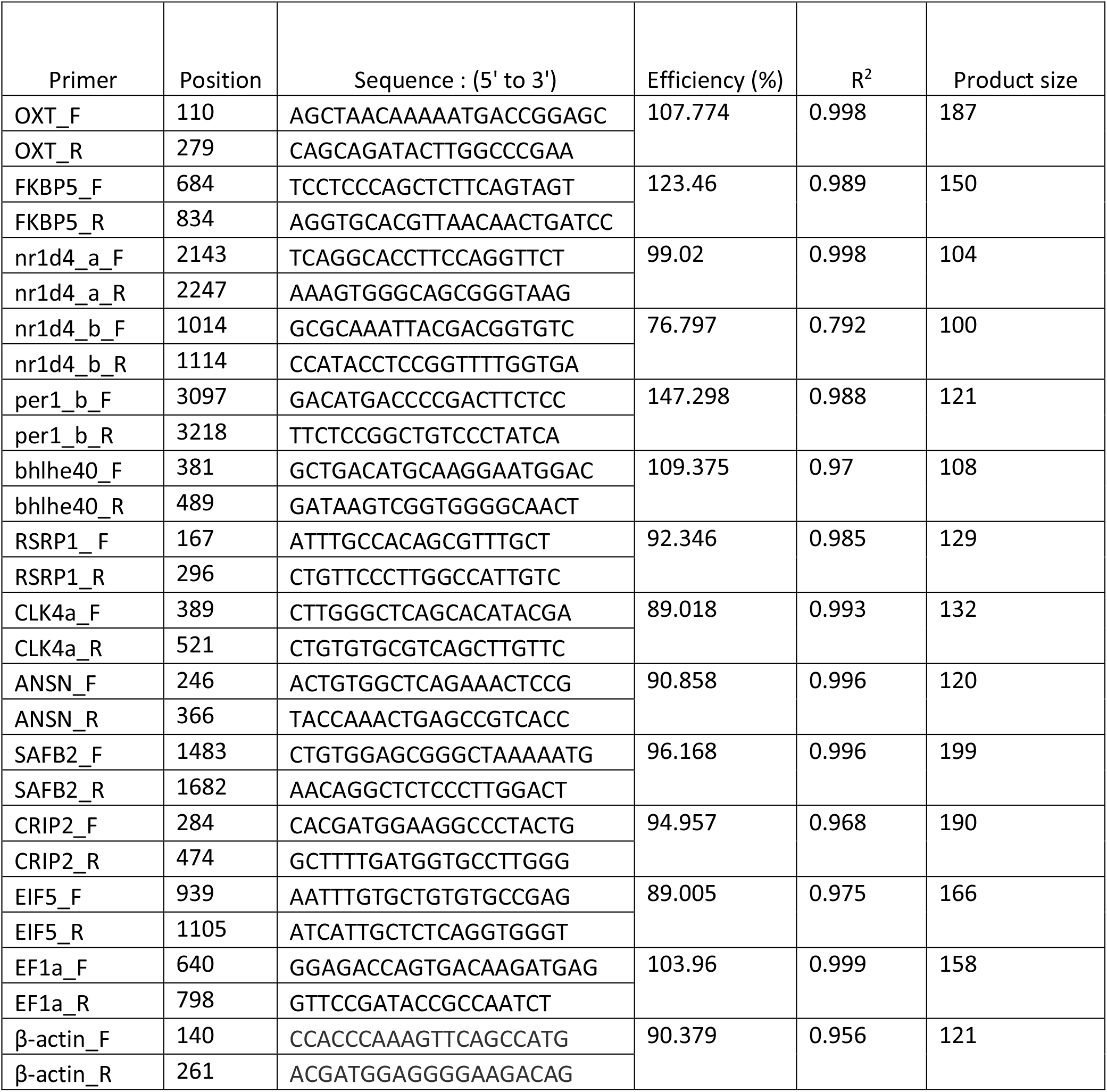
Oligos used in the current study.

